# Genome-wide association study of offspring birth weight in 86,577 women highlights maternal genetic effects that are independent of fetal genetics

**DOI:** 10.1101/034207

**Authors:** Robin N Beaumont, Nicole M Warrington, Alana Cavadino, Jessica Tyrrell, Michael Nodzenski, Momoko Horikoshi, Frank Geller, Ronny Myhre, Rebecca C. Richmond, Lavinia Paternoster, Jonathan P. Bradfield, Eskil Kreiner-Møller, Ville Huikari, Sarah Metrustry, Kathryn L. Lunetta, Jodie N. Painter, Jouke-Jan Hottenga, Catherine Allard, Sheila J. Barton, Ana Espinosa, Julie A. Marsh, Catherine Potter, Ge Zhang, Wei Ang, Diane J. Berry, Luigi Bouchard, Shikta Das, Early Growth Genetics (EGG) Consortium, Hakon Hakonarson, Jani Heikkinen, Øyvind Helgeland, Berthold Hocher, Albert Hofman, Hazel M. Inskip, Samuel E Jones, Manolis Kogevinas, Penelope A. Lind, Letizia Marullo, Sarah E. Medland, Anna Murray, Jeffrey C. Murray, Pål R. Njølstad, Ellen A. Nohr, Christoph Reichetzeder, Susan M. Ring, Katherine S Ruth, Loreto Santa-Marina, Denise M. Scholtens, Sylvain Sebert, Verena Sengpiel, Marcus A Tuke, Marc Vaudel, Michael N Weedon, Gonneke Willemsen, Andrew R Wood, Hanieh Yaghootkar, Louis J. Muglia, Meike Bartels, Caroline L. Relton, Craig E. Pennell, Leda Chatzi, Xavier Estivill, John W. Holloway, Dorret I. Boomsma, Grant W. Montgomery, Joanne M. Murabito, Tim D. Spector, Christine Power, Marjo-Ritta Järvelin, Hans Bisgaard, Struan F.A. Grant, Thorkild I.A. Sørensen, Vincent W. Jaddoe, Bo Jacobsson, Mads Melbye, Mark I. McCarthy, Andrew T. Hattersley, M. Geoffrey Hayes, Timothy M. Frayling, Marie-France Hivert, Janine F. Felix, Elina Hyppönen, William L. Lowe, David M Evans, Debbie A. Lawlor, Bjarke Feenstra, Rachel M. Freathy

**Author notes:** These authors contributed equally to this work (Robin N Beaumont, Nicole M Warrington). These authors jointly directed this work (David M Evans, Debbie A. Lawlor, Bjarke Feenstra, Rachel M. Freathy). A full list of consortium members is included in the **Supplementary Material** (Early Growth Genetics (EGG) Consortium). Corresponding authors: Dr. Rachel M. Freathy, University of Exeter, Medical School, Royal Devon and Exeter Hospital, Barrack Road, Exeter, UK, Prof. Debbie A. Lawlor, MRC Integrative Epidemiology Unit at the University of Bristol, Oakfield House, Oakfield Road, Bristol, UK, Dr. Bjarke Feenstra, Department of Epidemiology Research, Statens Serum Institut, Artillerivej 5, DK-2300 Copenhagen S, Denmark.

## Abstract

Genome-wide association studies (GWAS) of birth weight have focused on fetal genetics, while relatively little is known about how maternal genetic variation influences fetal growth. We aimed to identify maternal genetic variants associated with birth weight that could highlight potentially relevant maternal determinants of fetal growth.

We meta-analysed GWAS data on up to 8.7 million SNPs in up to 86,577 women of European descent from the Early Growth Genetics (EGG) Consortium and the UK Biobank. We used structural equation modelling (SEM) and analyses of mother-child pairs to quantify the separate maternal and fetal genetic effects.

Maternal SNPs at 10 loci (*MTNR1B, HMGA2, SH2B3, KCNAB1, L3MBTL3, GCK, EBF1, TCF7L2, ACTL9* and *CYP3A7*) showed evidence of association with offspring birth weight at *P*<5x10^-8^. The SEM analyses showed at least 7 of the 10 associations were consistent with effects of the maternal genotype acting via the intrauterine environment, rather than via effects of shared alleles with the fetus. Variants, or correlated proxies, at many of the loci had been previously associated with adult traits, including fasting glucose (*MTNR1B, GCK* and *TCF7L2*) and sex hormone levels (*CYP3A7*), and one (*EBF1*) with gestational duration.

The identified associations indicate effects of maternal glucose, cytochrome P450 activity and gestational duration, and potential effects of maternal blood pressure and immune function on fetal growth. Further characterization of these associations, for example in mechanistic and causal analyses, will enhance understanding of the potentially modifiable maternal determinants of fetal growth, with the goal of reducing the morbidity and mortality associated with low and high birth weights.

## Introduction

Individuals with birth weights approaching the lower and upper ends of the population distribution are more at risk of adverse neonatal and later-life health outcomes and mortality than those of average weight (1–5). The factors influencing birth weight involve both maternal and fetal genetic contributions in addition to the environment. Genome-wide association studies (GWAS) testing for common variant effects on *own* birth weight (“fetal” GWAS) have so far identified 60 robustly associated loci (6–8). The influence of common maternal genetic variation on offspring birth weight, beyond the effects of transmitted genetic variation, is poorly understood. Studies estimating the variance in birth weight explained by fetal or maternal genetic factors, using data on twins (9, 10), families (11) or mother-child pairs with genome-wide common variant data (8, 12), have consistently estimated a distinct maternal genetic contribution, which is smaller than the fetal genetic contribution, with estimates ranging from 3% to 22% of the variance explained (relative to 24% to 69% for fetal genetics).

Maternal genotypes may influence key maternal phenotypes, such as circulating levels of glucose and other metabolic factors, which could cross the placenta and affect the growth of the fetus. For example, women with hyperglycemia due to rare heterozygous mutations in the *GCK* gene have babies who are heavier at birth (provided the babies do not inherit the mutation) due to intrauterine exposure to high maternal glucose levels (13). Additionally, maternal genotypes may act upon other maternal attributes, such as vascular function or placental transfer of nutrients, which are also likely to influence fetal growth. Such maternal environmental effects could in turn influence fetal growth separately from the effects of any growth-related genetic variants that are inherited by the fetus directly from the mother (Figure 1). Supporting evidence for such effects from analyses of common genetic variants includes positive associations between maternal weighted allele scores for BMI or fasting glucose and offspring birth weight, and an inverse association between a maternal weighted allele score for systolic blood pressure and offspring birth weight (14).

**Figure 1.**
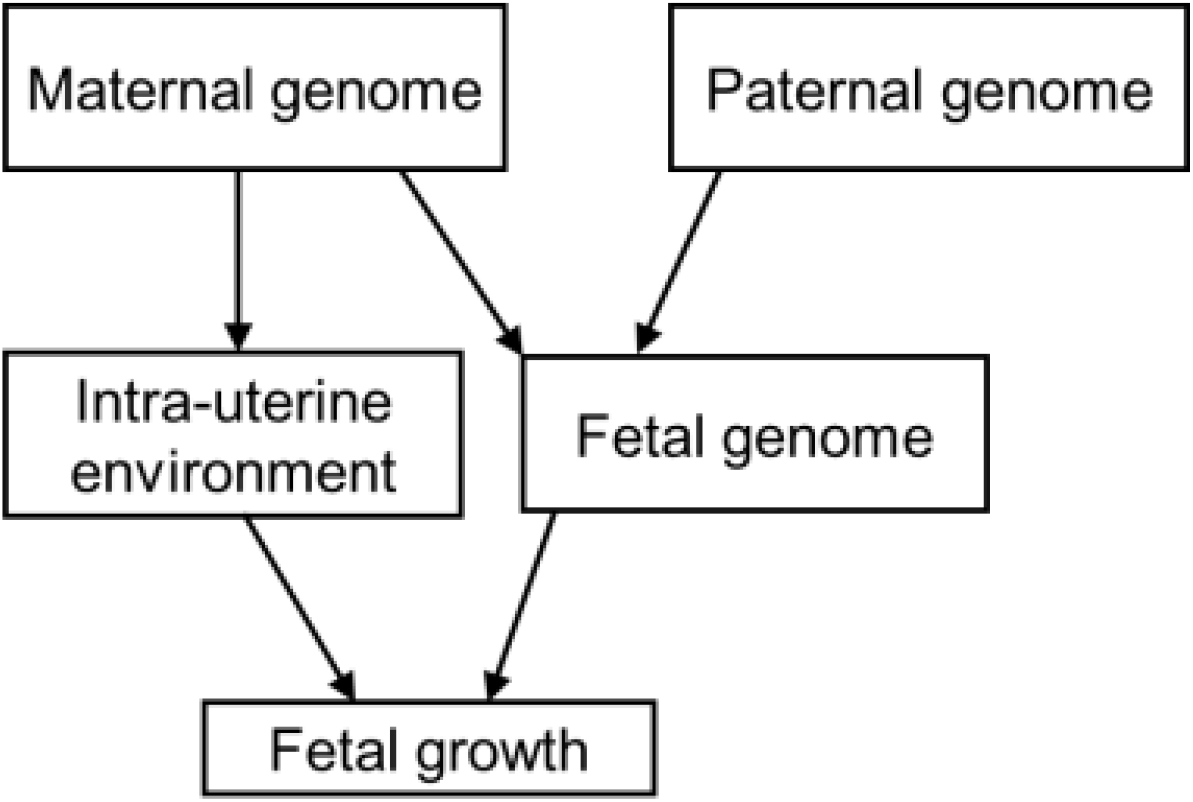
A schematic diagram illustrating that maternal genetic factors may influence fetal growth indirectly through the intra-uterine environment, or directly through inheritance by the fetus.

The goal of the current study was to apply a GWAS approach to identify *maternal* genetic variants associated with offspring birth weight. This could potentially highlight novel pathways by which the maternal genotype influences offspring birth weight through the intra-uterine environment. We performed a meta-analysis of GWASs of offspring birth weight using maternal genotypes in up to 86,577 women of European descent from 25 studies, including 37,945 participants from studies collaborating in the Early Growth Genetics (EGG) Consortium and 48,632 participants from the UK Biobank (**Supplementary Figure 1**).

## Results

The basic characteristics of study participants in the EGG Consortium discovery, EGG follow-up and UK Biobank GWAS analyses are presented in **Supplementary Tables 1, 2 and 3**, respectively.

### Maternal SNPs at 10 loci were associated with offspring birth weight at P < 5x10^-8^

We identified 10 autosomal loci that were associated with offspring birth weight at *P* < 5x10^-8^ (Figure 2; Table 1; **Supplementary Figures 2 and 3; Supplementary Table 4**). The linkage disequilibrium (LD) score regression intercept (15) from the overall meta-analysis was 1.009, so there was little change in the test statistics after adjusting for this inflation. Three of these loci (*KCNAB1, EBF1* and *CYP3A7*) were identified in UK Biobank data only, and the index SNPs were unavailable in the EGG Consortium data. Consideration of results for proxy SNPs at these three loci from the EGG meta-analysis is below in the next section. For the index SNPs at the other 7 loci, we observed no strong evidence of heterogeneity in allelic effects between the EGG Consortium and UK Biobank components of the meta-analysis (**Supplementary Figure 4; Supplementary Table 4**). The majority of the index SNPs mapped to non-coding sequence and were not in strong LD with any coding variants (r^2^<0.95), but the index SNP in *SH2B3*, rs3184504, is a non-synonymous coding variant (R262W). Approximate conditional analysis (see **Materials and Methods**) showed no evidence of secondary signals at any locus at *P* < 5x10^-8^. In combination, the 10 loci explained 1.4 % (SE = 1.2 %) of variance in birth weight, while the variance in birth weight captured by all autosomal genotyped variants on the UK Biobank array was considerably greater: 11.1 % (SE = 0.6 %).

**Figure 2.**
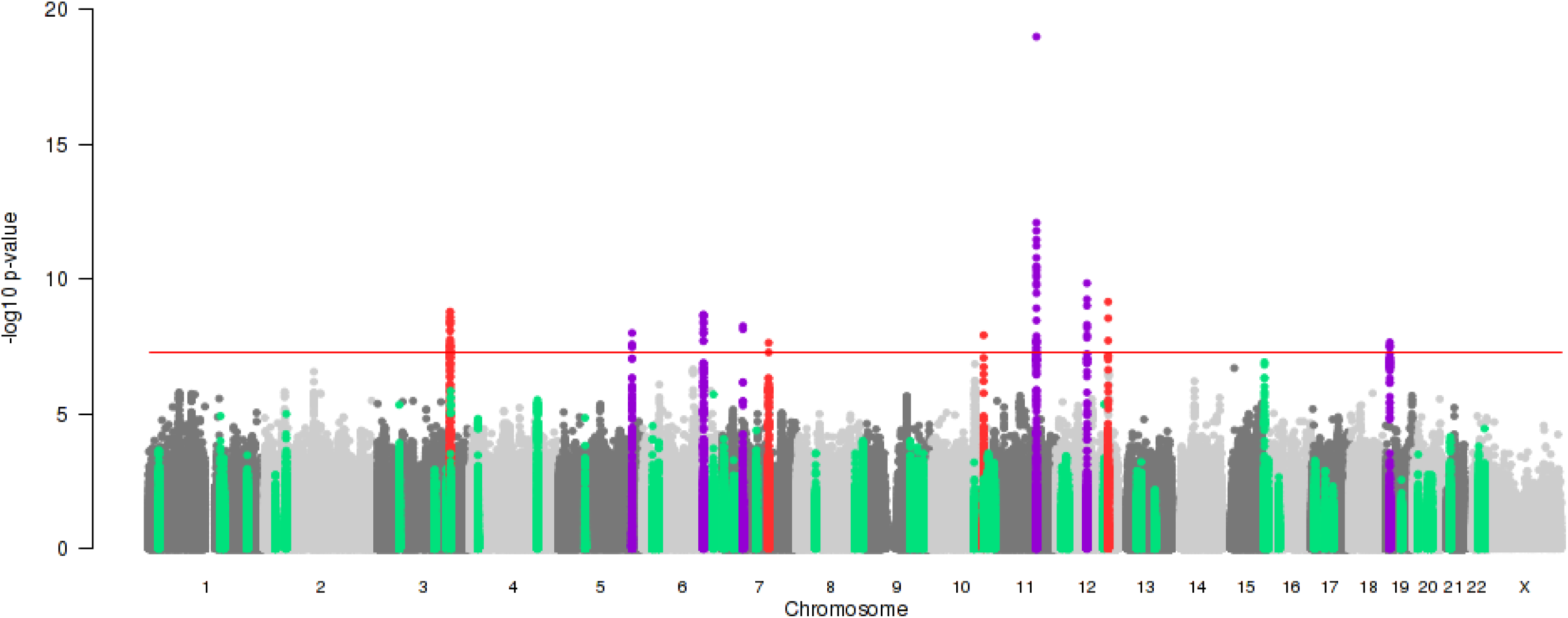
Manhattan plot of associations between 8,723,755 maternal autosomal SNPs and 17,352 maternal X-chromosome SNPs (all MAF >1%) and offspring birth weight from the meta-analysis of up to 86,577 women. SNP position across the chromosomes (x-axis) and results of association tests between maternal genotype and offspring birth weight adjusted for sex and, where available, gestational duration (-log10 P-value; left y-axis) are shown. The index maternal SNP from the current study and all SNPs within 500kb of that SNP are highlighted either in red, or in purple. Those in purple indicate loci at which the index maternal SNP from the current study was within 500k of an index SNP associated previously with own birth weight (i.e. in a “fetal GWAS of birth weight”) at *P* < 5x10^-8^ (8). SNPs within 500kb of the 54 other index SNPs previously identified in the fetal GWAS of birth weight are highlighted in green. The red, horizontal line indicates a *P* value of 5x10^-8^.

**Table 1.** Ten maternal genetic loci associated with offspring birth weight (*P* < 5 x 10^-8^) in a European ancestry meta-analysis of up to 86,577 women.

| *Locus* | Lead SNP | Chr | Position (bp, b37) | Alleles | EAF | EGG + UKBB meta-analysis |  |  |
| --- | --- | --- | --- | --- | --- | --- | --- | --- |
|  |  |  |  | Effect/Other |  | Effect (SE) | P-value | N |
| *MTNR1B* | rs10830963 | 11 | 92708710 | G/C | 0.29 | 0.052 (0.006) | $1.0 \times 10^{-19}$ | 71341 |
| *HMGA2* | rs1351394 | 12 | 66351826 | T/C | 0.49 | 0.034 (0.005) | $1.4 \times 10^{-10}$ | 68247 |
| *SH2B3* | rs3184504 | 12 | 111884608 | C/T | 0.52 | 0.033 (0.005) | $6.9 \times 10^{-10}$ | 68249 |
| *KCNAB1* * | rs7629460 | 3 | 155829938 | C/A | 0.59 | 0.039 (0.007) | $1.6 \times 10^{-9}$ | 48632 |
| *KCNAB1*† | rs9872556 | 3 | 155829855 | T/C | 0.57 | 0.029 (0.005) | $8.1 \times 10^{-8}$ | 68253 |
| *L3MBTL3* | rs9375694 | 6 | 130356608 | G/A | 0.31 | 0.035 (0.006) | $2.1 \times 10^{-9}$ | 68223 |
| *GCK* | rs2971669 | 7 | 44231778 | T/C | 0.23 | 0.038 (0.007) | $5.5 \times 10^{-9}$ | 68162 |
| *EBF1* * | rs12520982 | 5 | 157894747 | T/C | 0.74 | 0.041 (0.007) | $9.9 \times 10^{-9}$ | 48632 |
| *EBF1*† | rs2964484 | 5 | 157897437 | A/G | 0.71 | 0.031 (0.006) | $4.6 \times 10^{-7}$ | 67547 |
| *TCF7L2* | rs7903146 | 10 | 114758349 | T/C | 0.30 | 0.034 (0.006) | $1.2 \times 10^{-8}$ | 68253 |
| *ACTL9* | rs2918299 | 19 | 8787273 | C/T | 0.83 | 0.041 (0.007) | $2.2 \times 10^{-8}$ | 67603 |
| *CYP3A7* * | rs45446698 | 7 | 99332948 | G/T | 0.04 | 0.089 (0.016) | $2.4 \times 10^{-8}$ | 48632 |

Effects were aligned to the birth weight-raising allele and are presented in SD units (1 SD of birth weight = 484g (7)). Chr, chromosome; bp, base pair; b37, build 37; EAF, effect allele frequency; SE, standard error.
\*Index SNPs available in UK Biobank only.
†Results for proxy SNPs ( $r^2 = 1$ ) at *KCNAB1* and *EBF1* available in both UK Biobank and EGG (imputed to HapMap Phase 2) are shown below the index SNPs, with full details in **Supplementary Table 4** (in both cases there was weak evidence of heterogeneity of effect size between EGG and UK Biobank, $P < 0.01$ ). No proxy for rs45446698 at *CYP3A7* with $r^2 > 0.5$ was available in HapMap Phase 2.

### Birth weight-raising alleles at KCNAB1 and EBF1 were associated with longer gestational duration

The associations at *KCNAB1, EBF1* and *CYP3A7* resulted from analysis of UK Biobank data only, and index SNPs were unavailable in the EGG Consortium meta-analysis (imputed to HapMap Phase 2). To investigate further the evidence for association at these loci, we identified proxy SNPs (r^2^ = 1) for *KCNAB1* (rs9872556) and *EBF1* (rs2964484) that were available in HapMap Phase 2 (no proxy SNP was available at r^2^ > 0.5 at the *CYP3A7* locus). Meta-analysis of the EGG Consortium and UK Biobank data showed weaker evidence of association overall, with some evidence of heterogeneity between the EGG meta-analysis and UK Biobank (*P* = 0.008 and 0.007, respectively; Table 1; **Supplementary Table 4; Supplementary Figure 4**). In the UK Biobank, women reported the birth weight of their first child, but not the duration of gestation. In contrast, analyses of birth weight in all but one EGG study (QIMR, n=892) were adjusted for the duration of gestation. It is therefore possible that the associations observed with birth weight at *KCNAB1* and *EBF1* in the UK Biobank reflect primary associations with gestational duration. Look-ups of the index SNPs and HapMap 2 proxy SNPs in a published dataset of the top 10000 associated SNPs from a GWAS of gestational duration and preterm birth in 43,568 women (16) showed evidence of association at both *EBF1 (P* < 10^-12^) and *KCNAB1* (*P* < 10^-3^; **Supplementary Table 5**). The birth weight-raising alleles were associated with longer gestational duration.

### Five associated SNPs were independent of those identified in previous fetal GWAS of birth weight

The index SNPs at four of the identified loci (*SH2B3, KCNAB1, TCF7L2* and *CYP3A7*), mapped > 2 Mb away from, and were statistically independent of any index SNPs previously associated with birth weight at *P* < 5x10^-8^ in a fetal GWAS (r^2^ < 0.05) (8). A summary of candidate genes at these 4 loci is shown in **Supplementary Table 6** (corresponding information for the other loci was reported in (8)). At *MTNR1B* and *HMGA2*, the same index SNP was associated with birth weight in the same direction in both the current study and the previous fetal GWAS. At the four remaining loci, the maternal GWAS index SNPs were within 0.5 to 15 kb of previously reported fetal GWAS index SNPs with very different strengths of pairwise LD between the maternal and fetal GWAS index SNPs. At the *EBF1* and *ACTL9* loci the maternal and fetal GWAS index SNPs were in strong LD (r^2^ = 0.95 and 0.99, respectively), and the directions of association were consistent, suggesting that they were tagging the same causal variant. At the *L3MBTL3* locus the maternal and fetal directions of association were consistent, but the index SNPs were weakly correlated (r^2^ = 0.13), and conditional analyses in UK Biobank suggested weakening of the association when the *L3MBTL3* SNP identified in the fetal GWAS was accounted for (unconditional effect (SE) = 0.033 (0.007), *P*=1.4 x 10^-6^; conditional effect (SE) =0.024 (0.008), *P*=1.2 x 10^-3^; **Supplementary Table 7**). At the *GCK* locus the minor allele frequencies of the maternal and fetal GWAS index SNPs were very different (0.23 and 0.009, respectively), and in low pairwise LD (r^2^ = 0.002). Analysis conditional on the fetal GWAS index SNP in UK Biobank did not alter the association at the maternal index SNP (**Supplementary Table 7**), suggesting that at *GCK*, the maternal association with birth weight was distinct from the previously-reported fetal association.

### Structural equation modelling applied to UK Biobank data suggested most associations were driven by the maternal genotype

The partial overlap between associations identified in the current study and those identified in the previous fetal GWAS of birth weight (Figure 2; **Supplementary Figure 2**) illustrates the expected correlation between maternal and fetal genotypes (r ≈ 0.5). The associations between maternal genotype and birth weight identified here may represent indirect effects of maternal genotype on birth weight acting via the maternal intrauterine environment, or primary effects of the fetal genotype on birth weight that are captured (due to correlation) when assaying the maternal genotype, or a mixture of maternal and fetal effects. Analysis of UK Biobank data using structural equation modelling (SEM; n=78,674 male and female unrelated participants, of whom 33,238 individuals only reported their own birth weight, 20,963 women only reported the birth weight of their first child and 24,473 women reported their own birth weight and that of their first child); see **Materials and Methods**) provided estimates of maternal effects adjusted for fetal genotype, and vice versa, and suggested that the associations at the majority of the loci were driven by the maternal genotype (Figure 3; **Supplementary Table 8**). In particular, the adjusted maternal effects estimated at 7 of the loci (*MTNR1B, KCNAB1, GCK, EBF1, TCF7L2, ACTL9* and *CYP3A7*) were separated from the adjusted fetal effect estimates by at least 2 standard errors. The only locus at which the point estimate for the adjusted fetal effect was larger than that of adjusted maternal effect was *HMGA2*, suggesting this association was driven by the fetal genotype. Additional analyses (i) adjusting for fetal genotype in up to 8705 mother-child pairs, and (ii) comparing the unadjusted maternal effect estimates from the overall maternal GWAS (n=up to 86,577) with those from a published fetal GWAS (n=143,677), provided supporting evidence that the majority of the effects were maternally driven (**Supplementary Figure 5; Supplementary Tables 8 and 9**).

**Figure 3.**
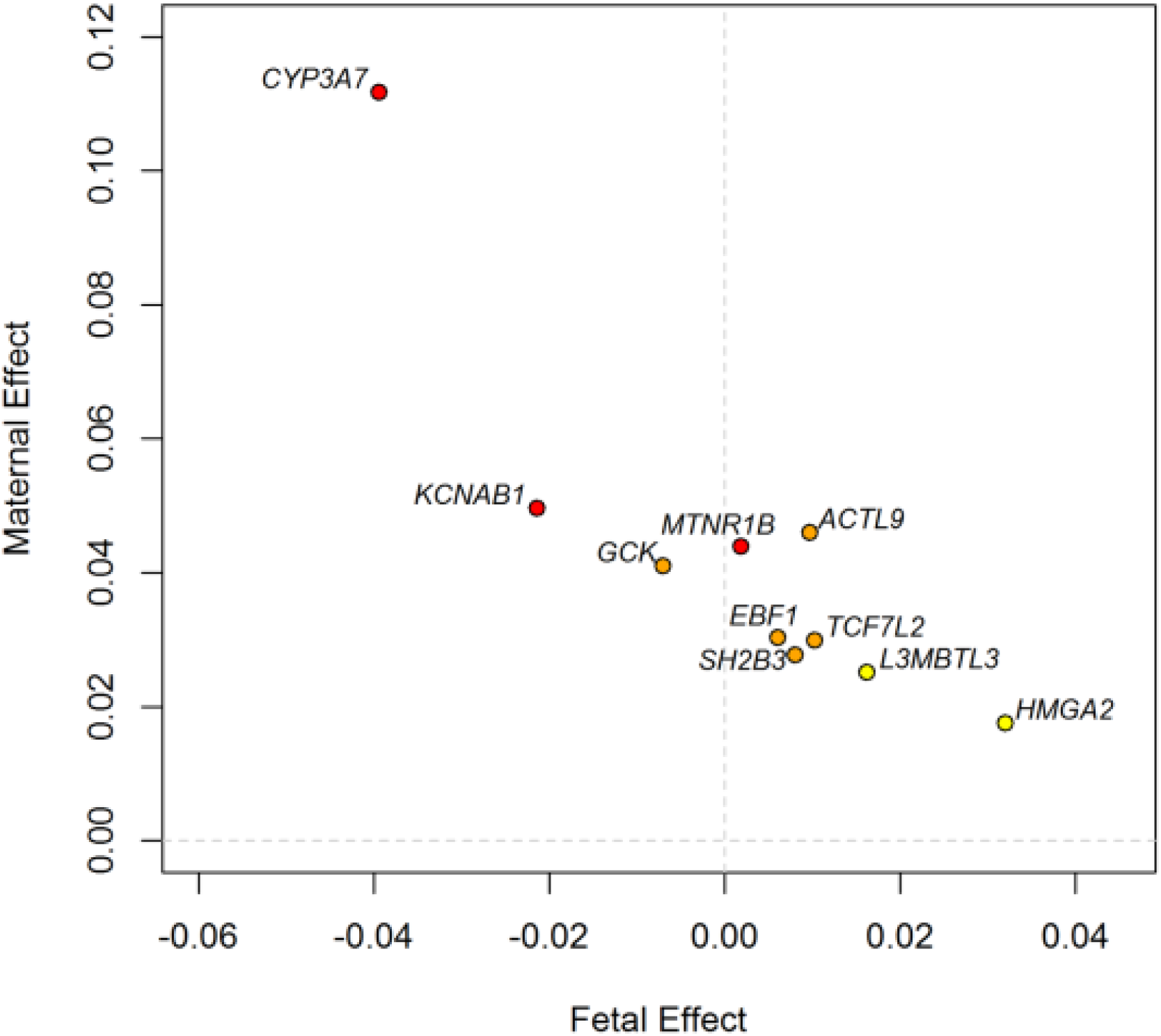
Independent maternal and fetal effects on birth weight at the 10 identified loci, estimated from a meta-analysis of results using a structural equation model (SEM) in unrelated UK Biobank participants with results using conditional analysis in maternal-fetal pairs. All SNPs are aligned to the birth weight-raising allele reported in Table 1. The colour of each dot indicates the maternal genetic association P-value for birth weight, adjusted for the fetal genetic association: red, *P* < 0.0001; orange, 0.0001 ≤ *P* < 0.001; yellow, 0.001 ≤ *P* < 0.05; white *P* > 0.05.

### Known associations at the identified loci highlighted potentially relevant maternal traits including fasting glucose, blood pressure, immune function and sex hormone levels

Look-ups of index SNPs (N=7 loci), or SNPs in close LD (N=2 loci at r^2^> 0.9; N=1 at r^2^=0.4), in available GWAS datasets for cardiometabolic and growth-related traits revealed several associations at *P <* 5x10^-8^ (**Supplementary Table 10**), while further information on previously-reported associations was obtained from the NHGRI-EBI catalog of GWAS (see **Materials and Methods**).

The maternal birth weight-associated variants at *MTNR1B, GCK* and *TCF7L2* loci are known to be associated with fasting glucose and Type 2 diabetes susceptibility (17, 18), with the glucose-raising allele associated with higher offspring birth weight.

The C-allele of the missense variant, rs3184504, in *SH2B3*, associated with higher birth weight in our study, has been associated with multiple cardiovascular traits (lower SBP and DBP (19, 20), altered lipid levels and lower risk of coronary artery disease, CAD (21, 22)), altered haematological traits (23–26), lower risk of autoimmune diseases and autoimmune disorders (27–32), lower kynurenine levels (33), higher risk of tonsillectomy (34) and higher risk of colorectal and endometrial cancer (35). The maternal birth weight-raising allele at *ACTL9* was in LD (r^2^ = 1) with alleles of nearby variants associated with lower risk of atopic dermatitis (36) and higher risk of tonsillectomy (34).

At the *CYP3A7* locus, a SNP in LD (rs34670419, r2=0.74) with our identified variant, rs45446698, has been associated with levels of the hormones, progesterone and dehydroepiandrosterone sulphate (DHEAS) (37). The maternal birth weight-raising allele was associated with lower hormone levels. The index SNP, rs12520982, at *EBF1* is in LD (r^2^ = 1) with rs2963463, recently associated with gestational duration and preterm birth (16). SNPs in weak LD (r^2^ < 0.1) with *EBF1* rs12520982 were previously associated with blood pressure traits (19, 38, 39), while rs12520982 showed moderate evidence of association with SBP (*P* = 1x10^-5^) and DBP (*P* = 1x10^-3^) in the UK Biobank data (**Supplementary Table 10**). There were no known prior associations with the birth weight-associated variation at *KCNAB1*.

The variants at *HMGA2* and *L3MBTL3* have been associated with adult height (40). At *HMGA2*, and possibly also at *L3MBTL3*, the association with birth weight is through the fetal allele, not the maternal allele, so these associations with adult height are relevant for offspring, not mother. Associations at *HMGA2* were additionally observed with other growth and development phenotypes: infant length (41); infant head circumference (42), and primary or permanent tooth eruption (43, 44).

To identify biological pathways underlying maternal regulation of birth weight, we performed gene-set enrichment analysis using MAGENTA (45). Seven pathways reached FDR <0.05, including three involved in the metabolism of xenobiotics (**Supplementary Table 11**).

## Discussion

In this study, we have identified variants in the maternal genome at 10 loci that are robustly associated with offspring birth weight. Five of the identified associations are independent of those reported in previous fetal GWAS of birth weight (8), bringing the total of known independent common variant associations with birth weight to 65. Since maternal and fetal genotype are correlated (r = 0.5), loci identified in GWAS of birth weight to date could either represent effects of the maternal genotype, acting via the intrauterine environment, or direct effects of the fetal genotype, or a mixture of the two (Figure 1). Our analyses, and those of 58 previously reported loci (46), suggest that while the majority of the 65 known associations indicate direct effects of the fetal genotype, at least 7 associations from the current study (those at *MTNR1B, EBF1, ACTL9, KCNAB1, GCK, TCF7L2* and *CYP3A7*, of which the first 3 were initially identified in fetal GWAS (8)) indicate maternal intrauterine effects.

The index SNP, rs45446698, at the *CYP3A7* locus, is an eQTL for *CYP3A7* in adrenal gland tissue (47). The *CYP3A7* gene is part of the cytochrome P450 family 3 subfamily A gene cluster, which encodes enzymes responsible for the metabolism of multiple and diverse endogenous and exogenous molecules (48), and SNP rs45446698 tags a haplotype of 7 highly correlated variants in the *CYP3A7* promoter, known as the *CYP3A7*1C* allele (49, 50). The *CYP3A7* gene is predominantly expressed in fetal development, but *CYP3A7*1C* results in expression in adult carriers (50, 51). The *CYP3A7*1C* allele, and correlated SNPs, have been associated with circulating levels of DHEAS, progesterone and 2-hydroxylation pathway estrogen metabolites (37, 52, 53). There were no associations between offspring birth weight and maternal SNPs at each of 9 loci (independent of *CYP3A7*) that are also known to influence levels of DHEAS or progesterone (37, 54) (data not shown), suggesting that neither DHEAS nor progesterone levels *per se* are likely to explain the association with birth weight. Since CYP3A enzymes metabolize a diverse range of substrates, there are many possible mechanisms by which maternal *CYP3A7*1C* might be associated with birth weight. In our conditional analysis, we observed weak evidence of an independent association with the fetal allele at this locus in the opposite direction to that of the maternal allele. Further analyses in larger samples will be required to confirm this and to investigate possible mechanisms underlying this association. However, the association at this locus, together with the results of the gene-set enrichment analysis, which highlighted pathways involved in xenobiotic metabolism, suggests that it is a key avenue for future research into fetal outcomes.

The birth weight-raising maternal alleles at the identified loci (*MTNR1B, GCK* and *TCF7L2*) are strongly associated with higher fasting glucose and Type 2 diabetes in non-pregnant adults (17, 18), and with glycemic traits and gestational diabetes mellitus in pregnant women (55–57). The association between raised maternal glucose and higher offspring birth weight is the result of higher fetal insulin secretion in response to increased placental transfer of glucose (58). Our results confirm previous maternal candidate gene associations with birth weight at *TCF7L2* and *GCK* (55, 59, 60) and demonstrate the key role of maternal glucose levels in influencing offspring birth weight (14, 61). Notably, the Type 2 diabetes risk allele at each of these three loci was not associated with birth weight independently of the maternal allele when present in the fetus. This is contrary to what has been seen at other Type 2 diabetes loci such as *ADCY5* and *CDKAL1*, where risk alleles in the fetus were associated with lower birth weight (8). However, there is an additional low frequency fetal variant at the *GCK* locus, which is independent of the glucose-raising maternal variant associated with higher birth weight in the current study (8). Taken together with the known effects on birth weight of both maternal and fetal rare heterozygous *GCK* mutations (13), a complex picture of allelic variation relevant to fetal growth is emerging at this locus.

The association with birth weight at *HMGA2* was previously identified in a fetal GWAS of birth weight (same index SNP) (8), and our analyses showed that the maternal SNP in our study was probably capturing a direct effect of the SNP in the fetus on skeletal growth, given previous associations with infant length, head circumference and adult height (40–42). The *L3MBTL3* locus identified in our study is also a known height locus, and the associated variant was correlated (r^2^ = 0.13) with a SNP associated with birth weight in the previous fetal GWAS (8). It was less clear from our analyses whether the association at *L3MBTL3* originated from the maternal or fetal genotype. However, analyses of maternal height alleles transmitted to offspring vs. those not transmitted to offspring suggest that the majority of the association between maternal height and offspring birth weight is due to direct effects of fetal inherited alleles (62).

Our exploration of known associations at the remaining 4 loci indicated a number of potentially relevant maternal traits that could influence birth weight via the intrauterine environment, including higher blood pressure (associations at *SH2B3* and suggestive associations at *EBF1*, both between the blood pressure raising maternal allele and lower offspring birth weight), which has been causally associated with lower birth weight in Mendelian randomization analyses (14), and immune function (associations at *SH2B3* and *ACTL9*). However, further studies are needed to elucidate the mechanisms at these loci and at *KCNAB1*, which showed no previous associations with other traits.

We observed weak evidence of heterogeneity of effect sizes between the EGG Consortium and UK Biobank components of our meta-analysis at the *KCNAB1* and *EBF1* loci, which led us to investigate possible explanations. A key difference was that birth weight was adjusted for duration of gestation in the majority of EGG studies, while the duration of gestation was unavailable in the UK Biobank. This raised the possibility that birth weight associations at *KCNAB1* and *EBF1* might arise from a primary effect on gestational duration, i.e. these loci could be primarily influencing the timing of delivery, rather than fetal growth. It is of course possible that the heterogeneity indicated false positive associations in the UK Biobank dataset that were not replicated in the EGG dataset. However, directionally consistent evidence of association with gestational duration and preterm birth in a recently published GWAS (*P* < 5x10^-8^ at *EBF1; P* < 10^-3^ at *KCNAB1*) suggests that this is unlikely.

There were some limitations to our study. First, the birth weight of first child was self-reported by mothers in the UK Biobank study, and so was likely subject to more error variation and potential bias than measured birth weight. However, maternal reports of offspring birth weight have been shown to be accurate (63, 64), and we showed that the birth weight of first child variable was associated with maternal smoking, height, BMI and socio-economic position in the expected directions. A second limitation of our study was that by performing a maternal GWAS of birth weight that does not account for the fetal genotype, the analysis was biased against identifying loci at which the fetal genotype exerts opposing effects. Proof-of-principle that such loci exist is demonstrated by the effects on birth weight of rare mutations in the *GCK* gene, which act in opposite directions when present in either mother or fetus, but result in normal birth weight if both mother and fetus inherit the mutation (13). Our analysis conditional on fetal genotype at the 10 loci using a novel method (46) had greatly increased power to resolve maternal vs. fetal effects compared with previous analyses in limited numbers of mother-child pairs (8). While it is not yet computationally feasible to run such an analysis genome-wide, future studies will benefit from considering maternal and fetal genotype simultaneously at the discovery stage and are thereby likely to uncover further loci.

In conclusion, we have identified 10 maternal genetic loci associated with offspring birth weight, 5 of which were not previously identified in fetal GWAS of birth weight, and at least 7 of which represent maternal intrauterine effects. Collectively, the identified associations highlight key roles for maternal glucose and cytochrome P450 activity and potential roles for maternal blood pressure and immune function. Future genetic, mechanistic and causal analyses will be required to characterize such intrauterine effects, leading to greater understanding of the maternal determinants of fetal growth, with the goal of reducing the morbidity and mortality associated with low and high birth weights.

## Materials and Methods

### Early Growth Genetics (EGG) Consortium discovery studies: genotyping, imputation and GWAS analysis

We studied 19,626 unrelated women of European ancestry from eleven studies with maternal genome-wide genotypes and offspring birth weight available. These included two sub-samples from the 1958 British birth cohort (1958BC-WTCCC2, n = 836; 1958BC-T1DGC, n = 858); the Avon Longitudinal Study of Parents and Children (ALSPAC, n = 7,304); a sub-sample of the Danish National Birth Cohort from the Genetics of Extreme Overweight in Young Adults study (DNBC-GOYA, n = 1,805); population-based controls from a case-control study of pre-term birth in the DNBC (DNBC-PTBCTRLS, n = 1,656); the Hyperglycemia and Adverse Pregnancy Outcome study (HAPO, n = 1,280); the Norwegian Mother and Child cohort study (MoBa, n = 650); the Northern Finland 1966 Birth Cohort study (NFBC1966, n = 2,035); the Netherlands Twin Register (NTR, n = 707); the Queensland Institute of Medical Research study of adult twins (QIMR, n = 892); the Twins UK study (TwinsUK, n = 1,603).

Genotypes in each study were obtained through high-density SNP arrays and up to ~2.5million autosomal SNPs were imputed to HapMap Phase II. Study protocol was approved at each study centre by the local ethics committee and written informed consent had been obtained from all participants and/or their parent(s) or legal guardians. Study descriptions and basic characteristics of samples in the discovery phase are presented in **Supplementary Table 1**.

Within each study, we converted offspring birth weight (BW, grams) to a z-score ((BW value - mean(BW))/ standard deviation(BW)) to allow comparison of data across studies. We excluded multiple births, stillbirths, congenital anomalies (where known), and births before 37 weeks of gestation (where known). We assessed the association between each SNP and offspring birth weight using linear regression of the birth weight z-score against maternal genotype (additive genetic model), with sex and gestational duration as covariables (gestational duration was unavailable in the QIMR study, which contributed 4.5 % of EGG participants). Ancestry principal components were included as covariables where necessary in the individual studies. Genome-wide association analyses were conducted using PLINK (65), SNPTEST (66), Mach2qtl (67) or Beagle (68) (see **Supplementary Table 1**).

### Genome-wide meta-analysis of 11 EGG Consortium discovery studies

Prior to meta-analysis, SNPs with a minor allele frequency (MAF) <0.01 and poorly imputed SNPs (info<0.8 (PLINK), r2hat <0.3 (MACH or Beagle) or proper_info <0.4 (SNPTEST)) were excluded. To adjust for inflation in test statistics generated in each cohort, genomic control (69) was applied once to each individual study (see **Supplementary Table 1** for λ values in each study). Data annotation, exchange and storage were facilitated by the SIMBioMS platform (70). Quality control of individual study results and fixed-effects inverse variance meta-analyses were undertaken by two meta-analysts in parallel at different study centres using the software package METAL (2009-10-10 release) (71). We obtained association statistics for a total of 2,422,657 SNPs in the meta-analysis for which at least 7 of the 11 studies were included. The genomic control inflation factor, λ, in the overall meta-analysis was 1.007.

### Follow-up of 18 SNPs in 13 additional EGG Consortium studies

We selected 15 SNPs that surpassed a P-value threshold of *P* < 1 x 10^-5^ for follow-up in additional, independent studies. Of these, one SNP (rs11020124) was in LD (r^2^ = 0.63, 1000 Genomes Pilot 1 data) with SNP rs10830963 at the *MTNR1B* locus known to be associated with fasting glucose and Type 2 diabetes (72). We assumed that these represented the same association signal. Given its robust association with maternal glycemic traits likely to impact on offspring birth weight, we took only rs10830963 forward for follow-up at this locus. We then used the National Human Genome Research Institute (NHGRI) catalog of published GWAS (http://www.genome.gov/gwastudies, accessed September 2012) to query the SNPs associated with birth weight between *P* = 10^-4^ and *P* = 10^-5^. We identified three further SNPs at loci with robust evidence (*P* < 5 x 10^-8^) of association with other phenotypes, and therefore higher prior odds of association with birth weight: rs2971669 near *GCK* (r^2^ = 0.73 with rs4607517 associated with fasting glucose)(60); rs204928 in *LMO1* (r^2^ = 0.90 with rs110419 associated with neuroblastoma)(73) and rs7972086 in *RAD51AP1* (r^2^ = 0.27 with rs2970818 associated with serum phosphorus concentration)(74). We took forward SNPs rs4607517, rs204928 and rs7972086 for follow-up at these loci, giving a total of 18 SNPs to be examined in additional studies.

The descriptions, genotyping details and basic phenotypic characteristics of the follow-up studies are presented in **Supplementary Table 2**. Of a total of thirteen follow up studies (n = 18,319 individuals), 9 studies (n = 15,288) provided custom genotyping of between 4 and 18 SNPs, while 4 studies (n = 3,031 individuals) had *in silico* genome-wide or exome-wide SNP genotypes available. Where SNPs were imputed, we included only those with quality scores (r2hat or proper_info) >0.8. We excluded directly genotyped SNPs showing evidence of deviation from Hardy-Weinberg Equilibrium at *P* <0.0028 (Bonferroni corrected for 18 tests). Where genotypes were unavailable for the index SNP, we used the SNP Annotation and Proxy (SNAP) Search Tool to find proxy SNPs based on LD in the 1000 Genomes Pilot 1 dataset (r^2^ > 0.8; https://www.broadinstitute.org/mpg/snap/ldsearch.php; accessed September 2012; see **Supplementary Table 12**).

### Preparation, quality control and genetic analysis in UK Biobank samples

UK Biobank data were available for 502,655 participants, of whom 273,463 were women (75), and of these women, 216,811 reported the birth weight of their first child (in pounds) either at the baseline or follow-up assessment visit. We converted pounds to kg (multiplying by 0.45) for use in our analyses. No information was available on gestational duration or offspring sex. A total of N=64,072 women with offspring birth weight data available also had genotype data available in the May 2015 data release. Women identified as not of British descent (N=9,681) were excluded from the analysis along with those reporting offspring birth weights of < 2.5 kg or > 4.5kg (N=5,479). ‘British-descent’ was defined as individuals who both self-identified as white British and were confirmed as ancestrally Caucasian using principal components analyses (http://biobank.ctsu.ox.ac.uk). A total of 1,976 of the women were asked to repeat the questionnaire at a follow-up assessment and therefore had two reports of birth weight of first child. Those with differing values were excluded (N=280). This resulted in N=48,632 women with both genotype data and a valid offspring birth weight value, which was z-score transformed for analysis (**Supplementary Table 3**). UK Biobank carried out stringent quality control of the GWAS genotype scaffold prior to imputation up to a reference panel of a combined 1000 Genomes Project Consortium and UK10K Project Consortium. We tested for association with birth weight of first child using a linear mixed model implemented in BOLT-LMM (76) to account for cryptic population structure and relatedness. Genotyping array was included as a binary covariate in the regression model. Total chip heritability (i.e. the variance explained by all autosomal polymorphic genotyped SNPs passing quality control) was calculated using restricted maximum likelihood (REML) implemented in BOLT-LMM (76). We additionally analysed the association between birth weight of first child and directly genotyped SNPs on the X chromosome in 45,445 unrelated women identified by UK Biobank as white British. We excluded SNPs with evidence of deviation from Hardy-Weinberg equilibrium (*P* < 1 x 10^-6^), MAF < 0.01 or overall missing rate > 0.015, resulting in 17,352 SNPs for analysis in PLINK v.1.07, with the first 5 ancestry principal components as covariates.

In both the full UK Biobank sample and our refined sample, birth weight of first child was associated with mother’s smoking status, maternal BMI and maternal height in the expected directions (**Supplementary Table 3**).

### Overall meta-analysis of discovery and follow-up samples

A flowchart of the overall study design is presented in **Supplementary Figure 1**. We performed inverse variance, fixed-effects meta-analysis of the association between each SNP and birth weight z-score in up to 25 discovery and follow-up studies combined (maximum total n = 86,577 women; 8,723,755 SNPs with MAF ≥ 0.01 plus 17,352 X-chromosome SNPs in 45,445 women) using METAL (71). To check for population substructure or relatedness that was not adequately accounted for in the analysis, we examined the intercept value from univariate linkage-disequilibrium score regression (15).

### Approximate conditional analysis

At each of the identified loci we looked for the presence of multiple distinct association signals in the region 1Mb up- and down-stream from the lead SNP through approximate conditional analysis. Conditional and joint analysis (COJO) in the analysis program, Genome-wide Complex Trait Analysis (GCTA) (77) was applied to identify secondary signals that attained genome-wide significance (*P* < 5 x 10^-8^) using a sample of 10,000 individuals selected at random from the UK Biobank to approximate patterns of LD between variants in these regions.

### Candidate gene search

To search for candidate genes at the 4 loci not already covered by the previous fetal GWAS of birth weight (8), we identified the nearest gene, searched PubMed for relevant information on genes within 300kb of the index SNP, and queried the index SNP for eQTL or proxy SNPs (r^2^ > 0.8) reported from GTEx v4, GEUVADIS, and 11 other studies using Haploreg v4.1 (http://archive.broadinstitute.org/mammals/haploreg/haploreg.php).

### Estimating maternal and fetal genetic effects at the identified loci

The full details of the structural equation modelling (SEM) method for estimating the conditional fetal and maternal effects are described elsewhere (46). Briefly, we fitted a structural equation model to three observed variables from the UK Biobank study; the participant’s own self-reported birth weight, the birth weight of the first child reported by the women and the genotype of the participants. Our model included two latent variables; one for the individual’s mother (i.e., grand-maternal genotype) and one for the genotype of the participant’s offspring. We know these latent variables are correlated on average 50% with the individual’s own genotype, hence the path coefficient between each of the latent variables and the observed genotype was set to 0.5. Our model also included residual error terms for the participant’s own birth weight and the birth weight of their first child, a covariance parameter to quantify similarity between the error terms, and a variance parameter to model variation in the observed genotype. Using this model, we were able to simultaneously estimate the effect of maternal and fetal genotypes on offspring birthweight To fit the SEM, we used OpenMx (78) in R (version 3.3.2) (79) with the raw UK Biobank data, and the P-value for the fetal and maternal paths was calculated using a Wald test. We fitted a second SEM without the child and maternal path to conduct a 2 degree of freedom test for the effect of the SNP on birth weight.

Genotype data from the UK Biobank May 2015 release was used for analysis. We included 57,711 participants who reported their own birth weight and 45,436 women who reported the birth weight of their first child, giving a total of a total of 78,674 unique individuals in the analysis (24,473 women had both their own and their offspring’s birth weight). Individuals who were non-European or were related to others in the sample, or who were part of multiple births, were excluded. The birth weight of offspring phenotype was prepared as described above, while own birth weight was prepared as described previously (8). We adjusted the individuals’ own birth weight for sex, and both birth weight measures for the 12 genetically determined principal components and genotyping batch before creating z-scores for analysis.

We analysed up to 8705 mother-child pairs from 4 studies with both maternal and fetal genotypes available (ALSPAC, EFSOCH, HAPO (non-GWAS) and DNBC-PTBCTRLS). We used linear regression to test the association between birth weight z-score and maternal genotype conditional on fetal genotype and vice versa (also adjusting analyses for sex and gestational duration). We combined the results from the individual studies using inverse variance meta-analysis with fixed effects. We performed a further meta-analysis to combine the overall estimates with those from the SEM using UK Biobank data.

### Look-ups in published GWAS and NHGRI GWAS catalog

We looked up associations between the 10 identified loci and various anthropometric and cardiometabolic traits in available GWAS result sets. The traits and sources are shown in **Supplementary Table 10**. Where the index SNPs at *KCNAB1, EBF1* and *CYP3A7* were unavailable, we used proxies (r^2^ = 0.99, 1.00 and 0.41, respectively). Since GWAS summary statistics for blood pressure were not publicly available, we used the UK Biobank May 2015 genetic data release and tested associations between the SNPs and systolic and diastolic blood pressure (SBP and DBP) in 127,968 and 127,776 British descent participants, respectively. Two blood pressure readings were taken approximately 5 minutes apart using an automated Omron blood pressure monitor. Two valid measurements were available for most participants, and the average was taken. Individuals were excluded if the two readings differed by more than 4.56SD, and blood pressure measurements more than 4.56SD away from the mean were excluded. We accounted for blood pressure medication use by adding 15 mmHg to the SBP measure and 10 mmHg to the DBP measure in those reporting regular use of any antihypertensive. Blood pressure was adjusted for age, sex and centre location and then inverse normalized before analysis.

We additionally queried the NHGRI-EBI catalog of published GWAS (http://www.ebi.ac.uk/gwas/home, last accessed 2 August 2017) for associations P < 5x10^-8^ between any additional traits or diseases and SNPs within 500kb of, and in LD with, the index SNP at each locus.

### Gene set enrichment analysis

We used Meta-Analysis Gene-set EnrichmeNT of variant Associations (MAGENTA) to test for pathway-based associations using summary statistics from the overall meta-analysis (45). The software mapped each gene to the SNP with the lowest P value within a 110kb upstream and 40kb downstream window. The P value (representing a gene score) was corrected for confounding factors such as gene size, SNP density and LD-related properties in a regression model. Genes within the HLA-region were excluded. Genes were then ranked by their adjusted gene scores. The observed number of gene scores in a given pathway with a ranked score above a given threshold (95^th^ and 75^th^percentiles) was calculated and this statistic was compared with 1,000,000 randomly permuted pathways of the same size. This generated an empirical *P* value for each pathway, and we considered pathways reaching FDR < 0.05 to be of interest. The 3230 biological pathways tested were from the BIOCARTA, Gene Ontology, Ingenuity, KEGG, PANTHER and REACTOME databases, with a small number of additional custom pathways.

## Acknowledgements

We are extremely grateful to the participants and families who contributed to all of the studies and the teams of investigators involved in each one. These include interviewers, computer and laboratory technicians, clerical workers, research scientists, volunteers, managers, receptionists and nurses. This research has been conducted using the UK Biobank Resource (Application numbers 7036 and 12703). For additional study-specific acknowledgements, please see Supplementary Material.

## Author contributions

Study Design (of individual contributing EGG Consortium studies): D.A.L., H.B., M-F. H., J.F.F., D.I.B., J.P.B., S.F.A.G., H.H., W.L.L., M.G.H., H.M.I., T.I.A.S., X.E., L. S-M., M.K., L.C., S.S., B.J., J.M.M., M.M., J.C.M., C.E.P., T.D.S., B.H., C.L.R., S.E., E.H., V.W.V.J., C.Power, M-R.J., A.T.H.

Sample Collection: D.A.L., S.M.R., H.B., E.K-M., B.F., F.G., M.M., M-F.H., V.W.V.J., M.B., J.P.B., H.H., S.F.A.G., J.W.H., H.M.I., B.H., C.R., C.L.R., T.I.A.S, E.A.N., G.W., L.S-M., B.J., M.K., L.C., S.D., G.W.M., M-R.J., C.Power, E.H., T.D.S., C.E.P, A.T.H.

Genotyping: D.A.L., D.M.E., S.M.R., M-R.J., J.C.M., L.B., J.F.F., J-J.H., H.H., S.F.A.G., J.W.H., M.I.M., L.M., C.L.R., C.Potter, L.P., X.E., M.V., Ø.H., P.R.N., G.W.M., C.E.P., T.D.S., T.M.F, R.M.F.

Statistical Analysis: R.N.B., N.M.W., J.T., R.M.F., R.C.R., D.A.L., D.M.E., E.K-M., L.P., B.F., F.G., C.A., J.F.F., J-J.H., G.Z., L.J.M., M.G.H., D.M.S., M.N., K.L.L., S.E.J., K.S.R., H.Y., A.R.W., M.A.T., A.M., M.N.W., M.H., C.L.R., C.Potter, A.E., S.D., V.H., J.N.P., S.E.M., P.A.L., A.C., D.J.B., R.M., V.S., J.A.M., W.A., S.M., S.J.B.

Writing and overall study direction: R.N.B., N.M.W., A.C., J.T., T.M.F., M-F.H., J.F.F., E.H., W.L.L., D.M.E., D.A.L., B.F. and R.M.F.

All authors reviewed and edited the manuscript.

